# Joint interactions with humans may pose a higher risk of zoonotic outbreaks than interactions with conspecifics among wildlife populations at human-wildlife interfaces

**DOI:** 10.1101/2021.07.19.452944

**Authors:** Krishna N. Balasubramaniam, Nalina Aiempichitkijkarn, Stefano S. K. Kaburu, Pascal R. Marty, Brianne A. Beisner, Eliza Bliss-Moreau, Malgorzata E. Arlet, Edward Atwill, Brenda McCowan

## Abstract

1. Pandemics caused by wildlife-origin pathogens, like COVID-19, highlight the importance of understanding the ecology of zoonotic transmission and outbreaks among wildlife populations at human-wildlife interfaces. To-date, the relative effects of human-wildlife and wildlife-wildlife interactions on the likelihood of such outbreaks remain unclear.
2. In this study, we used social network analysis and epidemiological Susceptible Infected Recovered (SIR) models, to track zoonotic outbreaks through wild animals’ joint propensities to engage in social-ecological co-interactions with humans, and their social grooming interactions with conspecifics.
3. We collected behavioral and demographic data on 10 groups of macaques (*Macaca* spp.) living in (peri)urban environments across Asia. Outbreak sizes predicted by the SIR models were related to structural features of the social networks, and particular properties of individual animals’ connectivity within those networks.
4. Outbreak sizes were larger when the first-infected animal was highly central, in both types of networks. Across host-species, particularly for rhesus and bonnet macaques, the effects of network centrality on outbreak sizes were stronger through macaques’ human co-interaction networks compared to grooming networks.
5. Our findings, independent of pathogen-transmissibility, suggest that wildlife populations in the Anthropocene are vulnerable to zoonosis more so due to their propensities to aggregate around anthropogenic factors than their gregariousness with conspecifics. Thus, the costs of zoonotic outbreaks may outweigh the potential/perceived benefits of jointly interacting with humans to procure anthropogenic food. From One Health perspectives, animals that consistently interact with both humans and conspecifics across time and space are useful targets for disease spillover assessments and control.

## Introduction

The COVID-19 pandemic has highlighted the importance of understanding infectious disease transmission among wildlife populations at human-wildlife interfaces (HWIs) (Gryseels et al. 2020; Townsend et al. 2020). Global population expansion has increased spatial overlap and contact rates between humans and wildlife (Dickman 2013; Nyhus 2016). The resultant HWIs are now widely recognized as ‘hotspots’ for the transmission and cross-species spillover of (anthropo)zoonotic (humans to wildlife, and vice-versa) infectious diseases (Cunningham 2017; Daszak et al. 2000). Despite this widespread recognition, there exists little quantitative, comparative research that unravels the *pathways* through which infectious agents may enter into and spread through wildlife populations at these locations. From an evolutionary perspective, such assessments can provide insights into how infectious disease risk influences, and is in-turn influenced by, (mal)adaptive responses in wildlife socioecology, behavioral flexibility, and risk-taking (McCabe et al. 2014; Silk et al. 2019). From a conservation and public health perspective, such assessments are critical to identify “edge” wildlife, that is individual animals or species ranging at HWIs which may transmit infectious agents into other wildlife and overlapping humans (Craft 2015; Engel & Jones-Engel 2011).

Research on disease transmission among wildlife populations at HWIs can be hampered by conceptual and methodological limitations. Traditional research on wildlife populations assumed that the probability of acquiring an infectious agent is equal across individuals within a defined area or cohort (Anderson & May 1992). In reality, wild animals at HWIs may interact with both other animals and humans, may do so to different extents across individuals, time, and space, and may form patterns of associations through such interactions that could influence zoonotic agent transmission. Social Network Analysis (SNA), through promising quantitative ways to evaluate animals’ tendencies to interact differently with different socio-ecological aspects of their environment (e.g., their conspecifics, other overlapping species including humans), offer exciting avenues to capture such associations and their impact on disease transmission (Drewe & Perkins 2015; Godfrey 2013; Silk et al. 2019). To-date, however, epidemiological studies that have implemented SNA have largely focused on animal-animal interactions, and often on single behavioral features that define such interactions (reviewed below). Some examples of wildlife-wildlife social networks that have been associated with increased risk of infectious agent transmission include shared use of space (e.g. Gidgee skinks, *Egernia stokesii*: Godfrey et al. 2009), contact associations (e.g., giraffes, *Giraffa camelopardalis*: VanderWaal et al. 2014a), aggression (e.g., meerkats, *Suricata suricatta*: Drewe 2010), and social grooming (e.g., Japanese macaques, *Macaca fuscata*: MacIntosh et al. 2012). Yet disease transmission among wildlife at HWIs may be driven by such multiple, potentially interplaying types of interactions, including inter-individual differences in animals’ interactions with conspecifics, humans, and anthropogenic features like contaminated water, soil, human foods, livestock, and other feral mammals (Balasubramaniam et al. 2020a; Bradley & Altizer 2007; Craft 2015). Among anthropogenically-impacted wildlife populations, it is therefore crucial to assess the relative effects of multiple (rather than single or specific types of) interactions – e.g. social interactions with conspecifics, co-occurrence or joint interactions with humans or other anthropogenic factors – on the risk of zoonotic transmission and resultant outbreaks.

Mathematical models offer critical insights into the occurrence of real-world epidemiological processes (Epstein & Axtell 1996). In this regard, network approaches have been extensively combined with bottom-up, compartmental ‘Susceptible Infected Recovered (SIR)’ models, that simulate disease spread by causing entities, which may be humans or other animals, to move across ‘susceptible’, ‘infected’, and ‘recovered’ disease states (Bansal et al. 2007; Brauer 2008). They do so at dynamic probabilities that, based on user specifications of model complexity, may depend on a combination of one or more pathogen-specific epidemiological variables (e.g., transmissibility, basic reproduction number: defined below), host contact patterns (e.g., spatial or social network connectedness), and host attributes (e.g., age-sex class) or intrinsic states (e.g., physiology, rates of recovery). To date, studies that have implemented SIR models in combination with wildlife spatial and social networks have revealed strong associations between network connectedness of the first-infected individual and simulated disease outcomes, such as pathogen extinction times (i.e. when all individuals have recovered from the disease and no more individuals can be infected) and outbreak sizes (mean % of infected individuals) (Carne et al. 2017; Rushmore et al. 2014; Sah et al. 2018). To-date, these models are yet to be implemented in the context of understanding the relative effects of anthropogenic factors and social behavior on the risk of zoonotic outbreaks in wildlife populations.

Human-nonhuman primate interfaces are well-suited to address the above gaps. Beyond sharing close evolutionary histories with humans (Hasegawa et al. 1985), several nonhuman primate (hereafter NHP) taxa have shared ecological niche space with humans for long periods of their evolutionary history (e.g., Chacma baboons, *Papio ursinus*, macaques, *Macaca* spp.), or following relatively recent exposure to human activities like ecotourism and habitat encroachment (e.g., chimpanzees, *Pan troglodytes*; mountain gorillas, *Gorilla gorilla beringei*) (reviewed in Fuentes & Hockings 2010; Lappan et al. 2020; Mckinney 2015). Unsurprisingly, human-primate interfaces are ‘hotspots’ for zoonotic transmission, spill-over, and emergence (Devaux et al. 2019; Kaur and Singh 2009; Lappan et al. 2020). NHPs may be vulnerable to many diseases contracted from humans (a recent study revealed that all African and Asian NHPs are vulnerable to infection from SARS-CoV-2: Melin et al. 2020), or act as natural reservoirs of pathogens that may invade and cause epidemics in otherwise uninfected human and wildlife populations. The genus *Macaca* are among the most ecologically and behaviorally flexible of all nonhuman primates. In the wild, many macaque species, particularly rhesus macaques, long-tailed macaques (*M. fascicularis*), and bonnet macaques (*M. radiata*), are considered ‘edge’ wildlife species that form ‘synanthropic’ associations (Klegarth 2017) with humans across a variety of anthropogenic landscapes (e.g. cities, temples, fields) where they experience highly spatiotemporally variant overlap and interactions with humans (Gumert 2011; Riley 2007). Influenced by their ecology and evolutionary history, macaques also show marked variation in social behavior with their conspecifics and (consequently) social networks (Balasubramanaim et al. 2018a; Thierry 2007). While (anthropo)zoonotic agents have been extensively documented among macaque populations that are synanthropic with humans (Balasubramaniam et al. 2020a), the social-ecological pathways that may underlie zoonotic transmission and outbreaks within such populations remain unclear.

Across human-macaque interfaces in India and Malaysia, we used network approaches combined with SIR models to evaluate the dynamics of zoonotic transmission and outbreaks among multiple groups and species of macaques. In doing so, we evaluated the relative vulnerability of these wildlife populations to zoonotic outbreaks through their social-ecological interactions with humans, and their social interactions with conspecifics. To capture patterns of macaques’ social-ecological interactions with humans, we constructed networks of macaques’ (nodes) shared tendencies to jointly engage in risk-taking or co-interacting with humans (edges), within the same time and location in the context of anthropogenic spaces (Balasubramaniam et al. 2021). To capture patterns of macaque-macaque social interactions, we constructed social ‘grooming networks’ that linked macaques based on the proportions of time they spent engaging in grooming their conspecifics. In a previous study, we revealed that macaques’ grooming relationships did not predict their tendencies to co-interact with humans, thereby establishing a premise to expect that their joint interactions with humans may offer different, somewhat independent pathways for zoonotic transmission than their social interactions with conspecifics (Balasubramaniam et al. 2021).

Independent of pathogen ‘transmissibility’ from an infected individual to a susceptible individual during its infectious period (Sah et al. 2018), we examined the impact of the behavioral ecology of wildlife host-species at HWIs on zoonotic outbreaks. Specifically, we examined the effects of hosts’ interaction- or network-type (social-ecological co-interactions with humans, versus grooming of conspecifics), host-species (rhesus, long-tailed, and bonnet macaque), and their interactions with the network connectedness or (hereafter) centrality of the first-infected macaque, on zoonotic transmission and outbreak sizes as predicted by epidemiological models. Consistent with previous research, we predicted that the connectedness or (hereafter) centrality of the first-infected macaque, irrespective of host-species and network-type, will be positively correlated to zoonotic outbreak sizes. We also examined whether the magnitude of this effect was different across network-type for each host-species, and across host-species for each network-type. Rhesus and long-tailed macaques, compared to bonnet macaques, typically show greater ecological flexibility and overlap with anthropogenic environments (Balasubramaniam et al. 2020b), as well as more nepotistic social systems characterized by greater tendencies for individuals to engage with specific subsets of group conspecifics than with others (Thierry 2007). Given these differences, across network-type for each host-species, we predicted that the co-interaction network centrality of first-infected macaques would have a stronger effect on outbreak sizes than grooming network centrality for rhesus macaques and long-tailed macaques, but that bonnet macaques would show the opposite effect. Across host-species for each network type, we predicted that the effect of co-interaction network centrality of first-infected macaques on outbreak sizes would be higher for rhesus macaques and long-tailed macaques compared to bonnet macaques, but that the effects of grooming network centrality on outbreak sizes would be the reverse (bonnet > rhesus and long-tailed macaques).

We also examined the effects of sociodemographic (sex, dominance rank) characteristics of the first-infected macaque on outbreak sizes. Since females and high-ranking individuals form the core of macaque grooming networks (Balasubramaniam et al. 2018a; Thierry 2007), we predicted that outbreak sizes through grooming networks would be higher when the first-infected individuals were females (versus males) and higher-ranking (versus lower-ranking) individuals. On the other hand, given the exploratory and increased risk-taking behavior of males resulting in their being more well-connected in co-interaction networks compared to females (Balasubramaniam et al. 2020b, 2021), we predicted that outbreak sizes through co-interaction networks would be higher when the first-infected individuals are males (versus females). Finally, we also explored whether the overall anthropogenic exposure of first-infected macaques, specifically their frequencies of interactions with humans, and time spent foraging on anthropogenic food, influenced zoonotic outbreak sizes through both network-types.

## Materials and Methods

### Study locations and subjects

We observed 10 macaque groups representing three different species at human-primate interfaces across three locations in Asia – four groups of rhesus macaques in Shimla in Northern India (31.05^0^N, 77.1^0^E) between July 2016 and February 2018, four groups of long-tailed macaques in Kuala Lumpur in Malaysia (3.3^0^N, 101^0^E) between September 2016 and February 2018, and two groups of bonnet macaques in Thenmala in Southern India (8.90^0^N, 77.10^0^E) between July 2017 and May 2018 (Supplementary Figure 1). All macaque groups were observed in (peri)urban environments, and their home-ranges overlapped with humans and anthropogenic settlements - e.g., Hindu temples (Shimla and Kuala Lumpur), recreational parks (outskirts of Kuala Lumpur, Thenmala), roadside areas (Thenmala, Shimla) – to varying extents (Balasubramaniam et al. 2020b; Kaburu et al. 2019; Marty et al. 2019a). Subjects were adult male and female macaques which were pre-identified during a two-month preliminary phase prior to data collection at each location. More details regarding the study locations, macaque group compositions and subjects, and observation efforts, may be found in our previous publications (Balasubramaniam et al. 2020b; Kaburu et al. 2019; Marty et al. 2019a) and in Supplementary Table 1.

### Data collection

We collected behavioral and demographic data in a non-invasive manner using observation protocols that were standardized across observers within and across locations (details in Balasubramaniam et al. 2020b, 2021). All data were collected for five days a week, between 9:00 am and 5:00 pm. To record and spatiotemporally capture variation in human-macaque social-ecological interactions for the construction of co-interaction networks, we used an *event sampling* procedure (Altmann 1974; Kaburu et al. 2018). For this we divided pre-identified parts of the home range of each macaque group in which human-macaque interactions were most likely to occur, into blocks of roughly equal area and observability. We visited these blocks in a pre-identified, randomized order each day. Within a 10-minute sampling period, we recorded interactions between any pre-identified subject macaque and one or more humans that occurred within that block, in a sequential manner. Human-macaque interactions included all contact and non-contact behaviors initiated by macaques towards humans (e.g., approach, aggression, begging for food), or vice-versa (e.g. approach, aggression, provisioning with food) within a three-meter radius of each other (more details in Kaburu et al. 2019).

To record macaques’ social behavior, and their overall anthropogenic exposure independent of spatiotemporal context, we used *focal animal sampling* (Altmann 1974). For this we followed individual subjects in a pre-determined, randomized sequence for 10-minute durations. In a continuous manner, we recorded, within each focal session, instances of social grooming, and dyadic agonistic interactions that involved aggression (threat, lunge, chase, attack) that was followed by submission (avoidance, silent bared teeth, flee), between the focal animal and its group conspecifics. We also recorded interactions between the focal animal and one or more humans in a continuous manner (see above for definitions). Once every two minutes, we ceased recording continuous data to conduct a *point-time scan* (Altmann 1974) of the focal animal’s main activity, i.e. one of resting, locomotion, socializing, interacting with a human, foraging on natural food, or foraging on anthropogenic food.

We entered all data into Samsung Galaxy Tablets using customized data forms created in HanDBase® application (DDH software). From these we exported and tabulated all the data into MS Excel and MS Access databases daily. All observers within and across locations passed inter-observer reliably tests using Cohen’s kappa (> 0.85) (Martin & Bateson 1993).

### Construction of co-interaction networks and grooming networks

From the human-macaque interactions collected using event sampling data, we constructed social-ecological co-interaction networks (Figure 1A). In these, nodes were individual macaques. Edges were based on the frequency with which pairs of macaques jointly engaged in interactions with humans at the same block and within the same event sampling session, per unit of event sampling observation time during which both members of the pair were present in the group and (thereby) observable (Balasubramaniam et al. 2021). We also constructed macaque-macaque social grooming networks using the focal sampling data (Figure 1B). In these, we linked individual macaques (nodes) based on the frequency which they engaged in social grooming interactions per unit of total focal observation times (edges) calculated for each pair of macaques during the period of their overlapping tenure in the group. Our use of different types of data (event sampling versus focal sampling) to construct co-interaction networks and social grooming networks respectively, minimized the potentially confounding effects of data inter-dependencies and sampling bias on our networks (Farine & Whitehead 2015).

**Figure 1:**
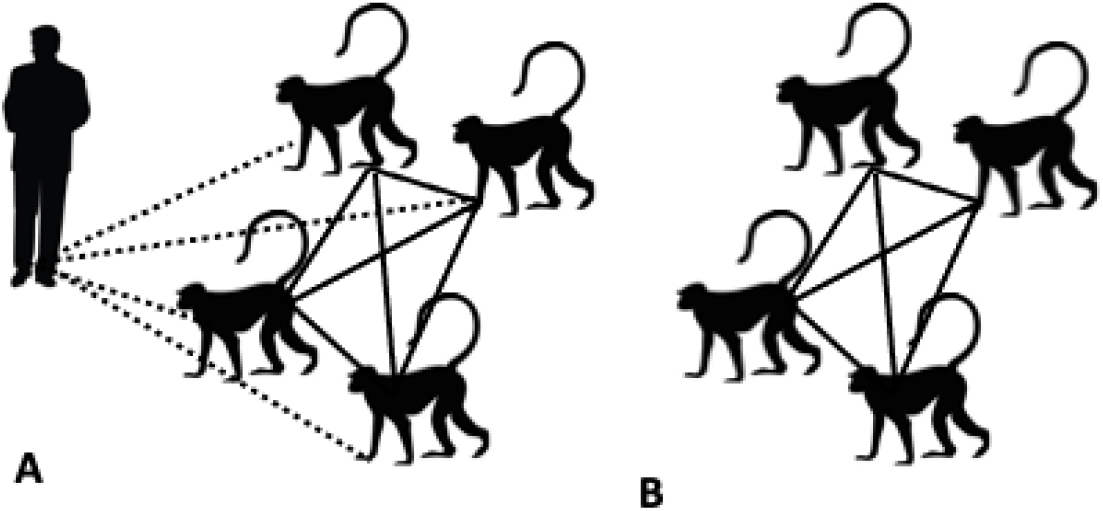
Construction of macaques’ (A) human co-interaction networks and (B) social grooming networks. Dotted lines represent macaques’ interactions with humans within the same (10-minute) time-window and space, which defined the edges of the co-interaction networks.

### Calculation of Network Measures

For each co-interaction network and grooming network, we calculated three measures of individual or node-level centrality. We calculated (1) weighted degree or strength centrality, i.e. the number and sum of the edge-weights of an individuals’ direct network connections, (2) betweenness centrality, i.e. the proportion of shortest paths connecting each pair of nodes that pass through a particular node, and (3) eigenvector centrality as the number and strength of an individuals’ direct and secondary network connections (reviewed in Farine & Whitehead 2015; Wey et al. 2008). These centrality measures were selected based on the decision-trees pertaining to choosing appropriate network measures provided by Sosa et al. (2020); they are among the most biologically relevant to modeling disease transmission pathways through animal networks (reviewed in Drewe & Perkins 2015). Specifically, strength indicates an individuals’ immediate susceptibility to acquiring infectious agents from infected conspecifics to whom they are directly connected. Betweenness indicates the tendency for an individual to function as a ‘bridge’ or a ‘conduit’ of disease spread. Eigenvector captures the reach of an individual within its network, and thereby its potential role in both acquiring and transmitting infectious agents to many other individuals. To account for cross-group differences in group size, we re-scaled centrality measures within each group into percentile values that ranged between 0 (least central individual) and 1 (most central individual).

### Macaque Sociodemographic Attributes and Overall Anthropogenic Exposure

From the data on dyadic agonistic interactions with clear winners and losers, we calculated macaques’ dominance ranks for each group, separately for male-male and female-female interactions, using the network-based Percolation and flow-conductance method (Package *Perc* in R: Fujii et al. 2016), a network-based ranking method that has been shown to yield animal rank orders that are highly consistent with those yielded by other, popularly used ranking methods in behavioral ecology, such as David’s score, I&SI ranks, and Elorating (Funkhouser et al. 2018). As with network centrality, we converted ordinal ranks of macaques within each group into percentile values that ranged between 0 (lowest-ranked individual) and 1 (highest-ranked individual). From the continuously collected focal sampling data, we calculated frequencies of human-macaque interactions per unit focal observation time. We also calculated, for each macaque, its time spent foraging on anthropogenic food as the ratio of the number of point-time scans in which it was foraging on anthropogenic food (Fa) to the total number of scans in which it was foraging on either anthropogenic food (Fa) or natural food (Fn), i.e. Fa/ (Fa + Fn).

### Zoonotic disease simulations

To simulate the spread of zoonotic agents of varying transmissibility (τ) on macaques’ co-interaction networks and grooming networks, we ran a series of Susceptible Infected Recovered (SIR) epidemiological models (using the *Epimdr* R package: Bjornstad 2020) (Figure 2A, B). We define ‘τ’ as a pathogen-specific characteristic, i.e. its probability of infecting a susceptible host within its infectious period which is a function of the probability of pathogenic infection 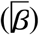 and recovery rate (γ), and is calculated as 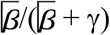 (Sah et al. 2018). For each network-type (human co-interaction, social grooming) and macaque group, we ran 5000 model simulations, 500 for each of 10 different values of τ ranging from 0.05 – 0.50 in increments of 0.05. These selections were based on the human literature that indicates that these values of τ correspond to zoonotic agents that range from low (e.g., influenza virus: Tuite et al. 2010), to moderate (e.g., respiratory pathogens like SARS-CoV-2: Arienzo & Coniglio 2020), to high (e.g., measles virus: Anderson & May 1992) contagiousness, and average basic reproduction numbers (R_0_) of between 1.6 – 14.0 (Rushmore et al. 2014; Sah et al. 2018). We thus ran a total of 100,000 simulations (5000 per macaque group times 10 groups times two network-types). In each simulation, we deemed all macaques within a group to be initially ‘susceptible’, and then infected one individual (node) at random with an artificial zoonotic agent of a given τ. A simulation proceeded using a discrete time, chain binomial method (Bailey 1957; Sah et al. 2018) that dynamically and temporally tracked the spread of infection through a weighted, undirected network through time (example in Figure 4B). In each simulation, animals were allowed to transition from ‘susceptible’ to ‘infected’ states, as a function of their network connections to individuals already in ‘infected’ states and the pathogen τ value. ‘Infected’ individuals were then allowed to transition into ‘recovered’ states at a fixed recovery rate (γ) of 0.2 that corresponds to an average infectious period of five days (Sah et al. 2018). Each simulation was allowed to proceed until the disease proceeded to extinction when there were no remaining infected individuals in the network. At the end of each simulation, we calculated the disease outcome of ‘mean outbreak size’, as the average % of infected macaques (the number of ‘infected’ individuals divided by the total number of individuals) across all time-units of the simulation. We also extracted, for each simulation, the identity of the first-infected macaque ‘k’ (Figure 2A) and calculated an average of zoonotic outbreak sizes from across all its first-infected simulation runs. We then matched this individual-level mean outbreak size with the sociodemographic characteristics, network centrality, and overall anthropogenic exposure of this (first-infected) individual.

**Figure 2:**
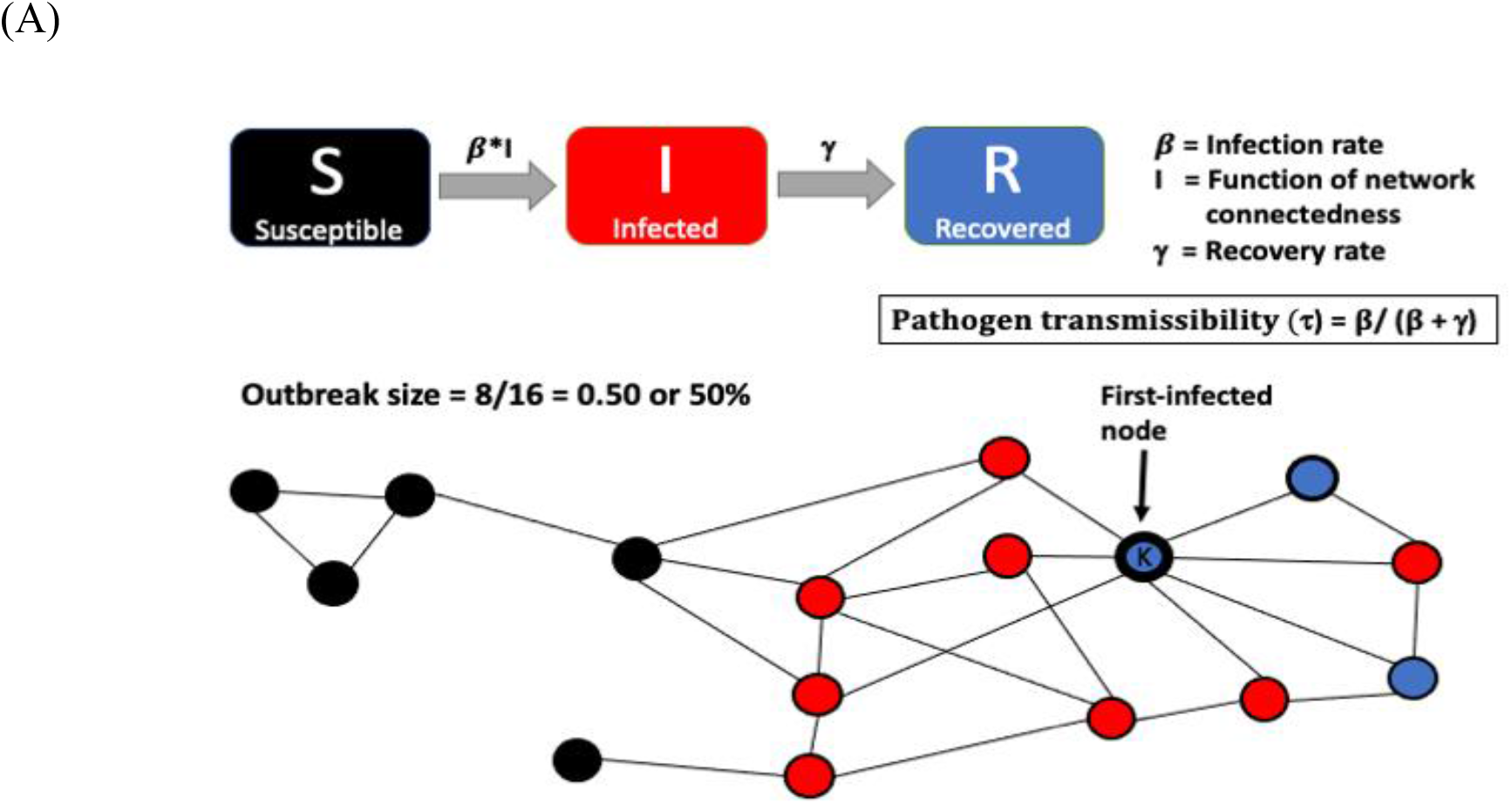
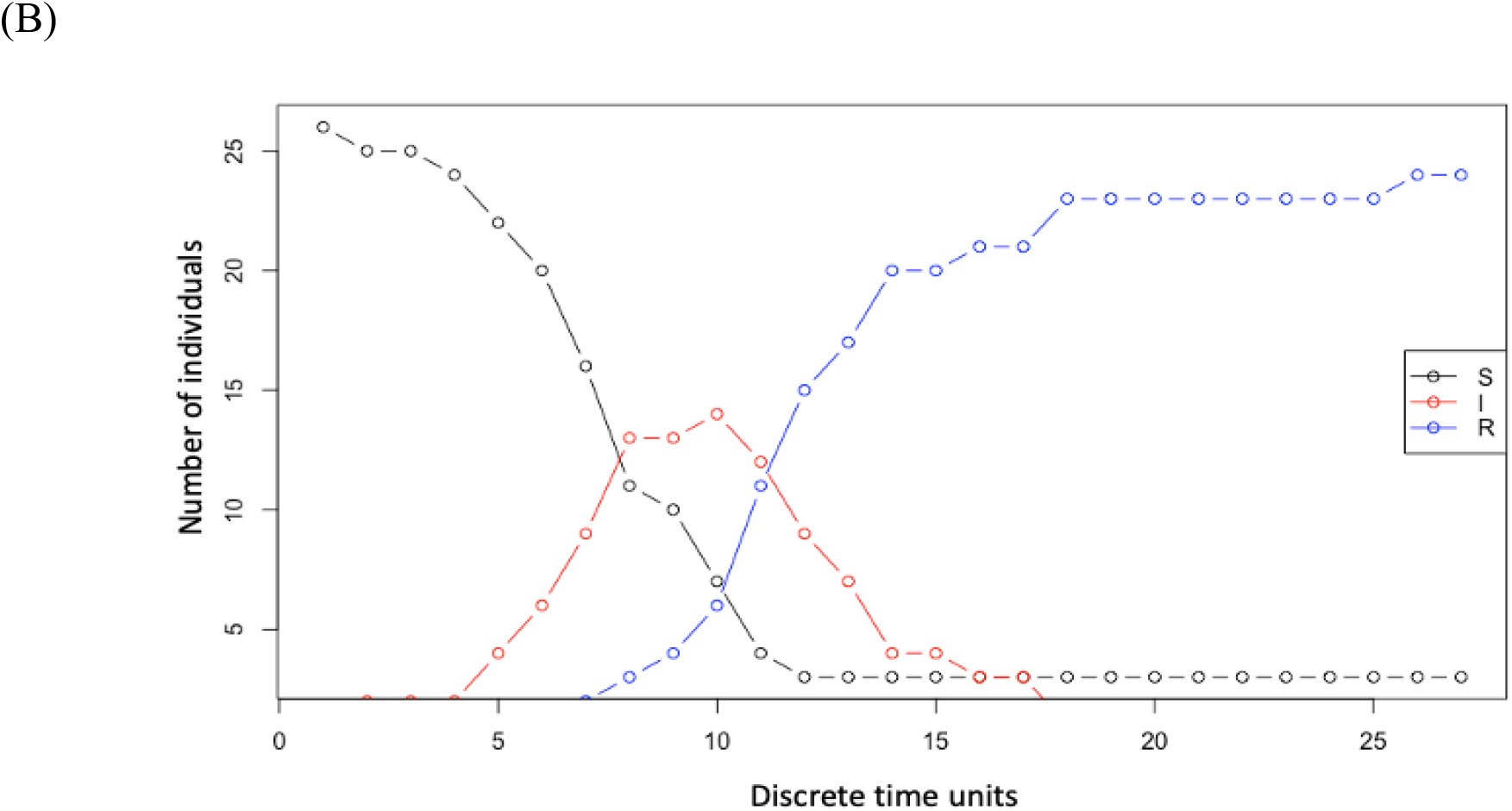
A typical Susceptible Infected Recovered (SIR) model simulation of network-mediated disease transmission (A), and an output from a single discrete time-based SIR model simulation (B).

**Figure 3:**
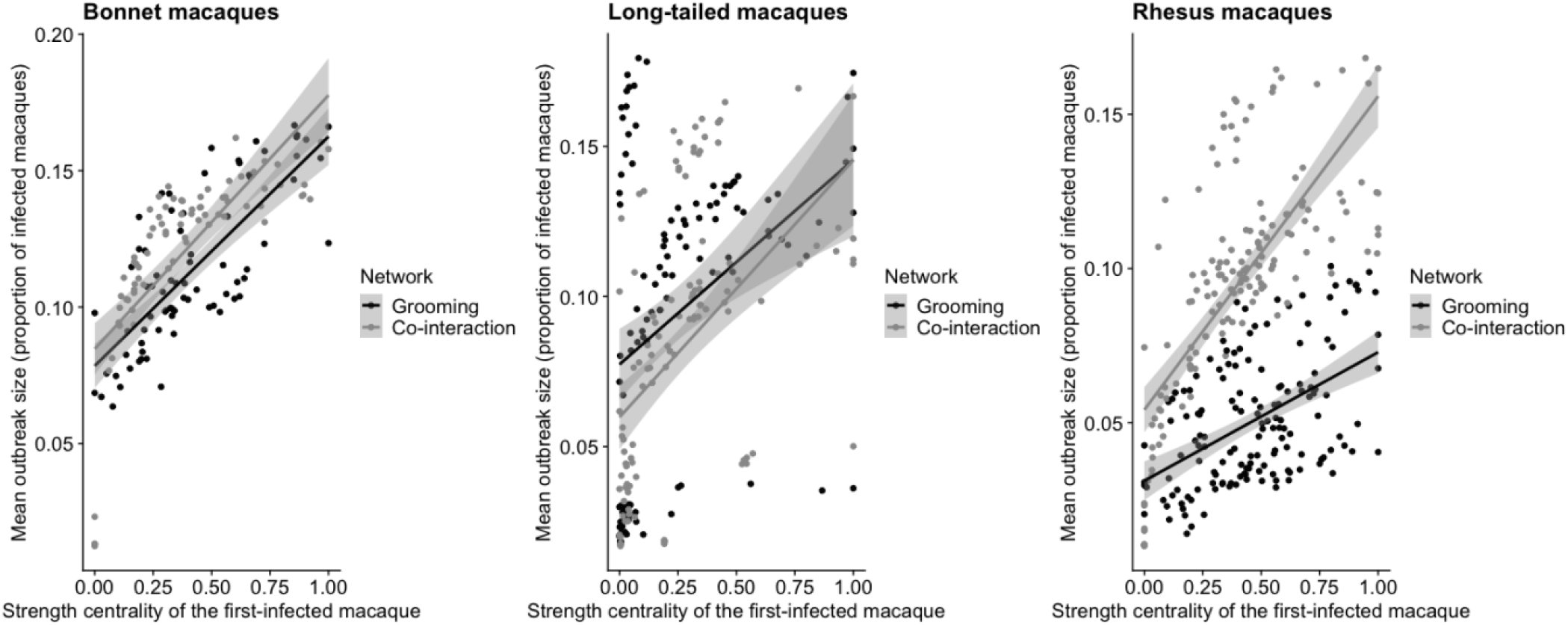
Scatterplots showing positive correlations between the strength centrality of first-infected macaques by network-type for each host-species.

**Figure 4:**
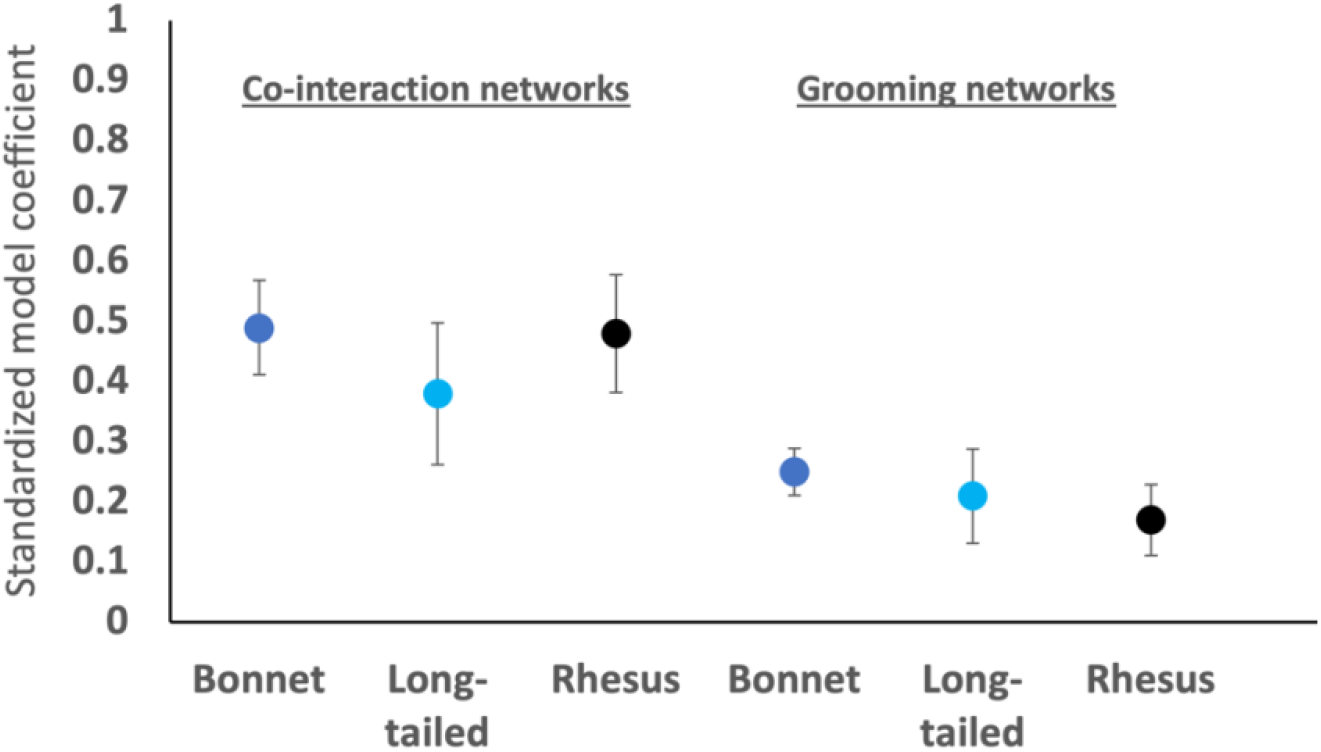
Plots of standardized model-coefficients (Y-axis; values from Table 2) to show the difference in effects of the strength centrality of first-infected macaques on zoonotic outbreak sizes through co-interaction networks and grooming networks. Coefficients for the same host-species are colored the same (dark blue = bonnet macaques; light blue = long-tailed macaques; black = rhesus macaques). Error bars represent 95% confidence intervals for each coefficient.

### Statistical Analysis

We used General Linear Mixed Models (GLMMs) implementing a corrected Akaike Information Criterion (AICc)-based model-selection criterion (packages *Lme4* (Bates et al. 2015) and *MuMIn* (Burnham et al. 2011) to test our predictions. In all GLMMs, we set mean outbreak size calculated at the level of the individual macaque through their co-interaction networks and/or grooming networks as the outcome variable. We used a Gaussian function since outcome variables did not deviate from a normal distribution (Shapiro Wilcoxon tests: p > 0.05 in each case). First, to examine the effect of the centrality of the first-infected macaque by network-type (co-interaction versus grooming) for a given host-species (bonnet or long-tailed or rhesus) on mean outbreak sizes, we ran three sets of three GLMMs each, one for each macaque species (details in Table 1A). In all models, we set the number of macaque subjects within the group (or ‘effective group size’) to be an offset variable, since group size can impact our outcome variable of mean outbreak sizes (Griffin & Nunn 2012). In all models, we also included ‘animal ID’ (a repeated measure for co-interaction networks and grooming networks) nested within macaque ‘group ID’ as a random effect to control for intraspecific variation. For each species, we ran three models, in each of which we included just one of the three different network measures of the centrality of the first-infected macaque, i.e. the strength, betweenness, or eigenvector, as a main effect. We used this approach in order to avoid the confounding effects of potential inter-dependencies of network centrality measures (Farine & Whitehead 2015). In each of these three models, we also included an interaction term of network centrality by network-type (co-interaction versus grooming), to determine whether the magnitude of these effects were different for different types of interactions. In all models, we also included, as main effects, the sociodemographic attributes (sex, dominance rank) and the overall anthropogenic exposure (frequencies of interactions with humans, proportions of time spent foraging on anthropogenic food) of the first-infected macaque. From each model-set of three models, we identified a single best-fit model with a difference in AICc of at least 8 points or lower than the next best-fit model (Burnham et al. 2011).

**Table 1:** Summary of GLMM sets to examine the impact of the centrality of the (A) the first-infected macaque by network-type for a given host-species, and (B) the first-infected macaque by species for a given network-type, on zoonotic outbreak sizes

| (A) Effects of the first-infected macaque by network-type for a given species |  |  |
| --- | --- | --- |
| Bonnet macaques | Long-tailed macaques | Rhesus macaques |
| 3 models | 3 models | 3 models |
| (on 76 individuals repeated across two network-types) | (on 112 individuals repeated across two network-types) | (on 151 individuals repeated across two network-types) |
| **(B) Effects of the first-infected macaque by species for a given network-type** |  |  |
| **Co-interaction networks** | **Grooming networks** |  |
| 3 models<br>(on 339 individuals across three species) | 3 models<br>(on 339 individuals across three species) |  |

Second, to examine the effect of the centrality of the first-infected macaque by species (bonnet versus long-tailed versus rhesus) for a given network-type (co-interaction or grooming) on mean outbreak sizes, we ran two sets of three GLMMs each, one for each network-type (details in Table 1B). Once again, we set the number of macaque subjects to be an offset variable, and included ‘group ID’ as a random effect, in all the models. For each network-type, we once again ran three models, in each of which we included just one of the three different measures of the centrality of the first-infected macaque as a main effect. In each of these three models, we also included an interaction term of network centrality by host-species (bonnet versus long-tailed versus rhesus), to determine whether the magnitude of these effects were different for different species. Again we included, as main effects, the sociodemographic attributes (sex, dominance rank) and the overall anthropogenic exposure (frequencies of interactions with humans, proportions of time spent foraging on anthropogenic food) of the first-infected macaque. From each model-set of three models, we identified a single best-fit model with a difference in AICc < 8 points from the next best-fit model (Burnham et al. 2011).

To account for inter-dependencies across network measures, we used a post-network ‘node-swapping’ randomization procedure to calculate permuted p (p_perm_) values for the observed model coefficients for predictor variables from each best-fit model (Farine & Whitehead 2015; Weiss et al. 2020). Specifically, we compared observed model coefficients to a distribution of coefficients generated by re-running the best-fit GLMM 1000 times, each following randomized re-assignments of the observed network centrality scores across individuals within each macaque group. All GLMMs met the necessary assumptions of model validity (i.e., distribution of residuals, residuals plotted against fitted values: Quinn & Keough 2002). All statistical tests were two-tailed, and we set the p values to attain statistical significance to be < 0.05.

## Results

### Impact of the centrality of first-infected individuals by network-type and host-species on zoonotic outbreak sizes

In support of our prediction, we found that across network-types and host-species, the strength centrality of the first-infected macaque, which better predicted outbreak sizes than betweenness centrality or eigenvector centrality (model 1 in Supplementary Tables 2A-C, 3A-B), was significantly, positively correlated to mean outbreak size (Tables 2 and 3; Figures 3 and 4). Moreover, the magnitude of these effects of first-infected macaque centrality on outbreak sizes varied across network types and species, although not always in the predicted directions.

**Table 2:** Standardized model coefficients from the best-fit GLMMs (model 1 from Supplementary Tables 2A, 2B and 2C) of network centrality of the first-infected ‘patient-zero’ macaque by network-type (co-interaction vs grooming) for a given host-species. In each model, we included macaque animal ID (repeated measure across network-type) nested within group ID as random effects, to account for intraspecific variation.

| Predictor | Model Coefficients |  |  |
| --- | --- | --- | --- |
|  | Bonnet macaques | Long-tailed macaques | Rhesus macaques |
| (Intercept) | 1.20* | 0.95* | 0.64* |
| Sex (males vs females) | -0.07* | -0.01 | -0.03 |
| Rank percentile | 0.01 | 0.05 | 0.04 |
| Network (grooming vs co-interaction) | -0.10** | 0.12** | -0.21** |
| Network strength (co-interaction) | 0.37** | 0.22** | 0.91** |
| Network strength (grooming) | 0.18** | 0.16** | 0.28** |
| Frequency of interactions with humans | 0.04 | 0.03 | 0.02 |
| Foraging on anthropogenic food | 0.04 | 0.01 | -0.04 |
| Network strength (grooming vs co-interaction) | -0.19** | -0.06 | -0.63** |

\*p < 0.05; \*\*p < 0.01 Note: p values were calculated after re-running the GLMMs using network centrality measures generated from 1000 post-network randomizations or node-swappings conducted on the co-interaction network and the grooming network.

<u>For a given host-species but across the two different types of networks</u>, we found a significant interaction between network-type and strength centrality for rhesus macaques and bonnet macaques, but not for long-tailed macaques (Table 2; Figure 3). As predicted, rhesus macaques showed a significantly stronger effect of the mean centrality of first-infected individuals on outbreak sizes through their co-interaction networks compared to their grooming networks (Table 2; Figure 3). In other words, disease-causing agents were likely to infect more individuals if they entered into a population by first infecting monkeys that were more central in human co-interaction networks, compared to by first infecting monkeys that were more central in grooming networks. Contrary to our predictions, bonnet macaques also showed the same (rather than the opposite) effect as rhesus macaques, although the magnitude of difference was somewhat lesser than for rhesus (Table 2; Figure 3). Finally, although the centrality of first-infected macaques within their co-interaction networks once again showed an overall greater effect on outbreak sizes than the centrality of macaques within their grooming networks for long-tailed macaques, this difference was not significant (Table 2; Figure 3). Moreover, long-tailed macaques also seemed to show separate groupings within each network-type (Figure 3), suggesting possible intra-specific differences in the effects of the network centrality of macaques within each network-type on outbreak sizes (see Discussion).

<u>For a given network-type but across host-species</u>, we found a significant interaction between species and strength centrality for both co-interaction networks and grooming networks (Table 3). For co-interaction networks, rhesus macaques showed the strongest effect of strength centrality on outbreak sizes as predicted. Contrary to our predictions, bonnet macaques fell within the range of rhesus macaques, and long-tailed macaques showed a significantly lower effect than both rhesus and bonnet macaques (Table 3; Figure 4). For grooming networks, the differences were in the directions we predicted – bonnet macaques showed the strongest effects of strength centrality on outbreak sizes, followed by long-tailed macaques, and finally rhesus macaques that showed a significantly lower effect compared to bonnet macaques (Table 3; Figure 4). For all three species, the magnitude of the effects of strength centrality on outbreak sizes was markedly greater for co-interaction networks compared to grooming networks (Figure 4). In other words, across host-species, the infection of macaques that were central in their co-interaction networks led to consistently higher zoonotic outbreaks (more individuals infected) than the infection of macaques that were central in their grooming networks.

**Table 3:** Standardized model coefficients from the best-fit GLMMs (model 1 in Supplementary Table 3A and 3B) of network centrality of the first-infected ‘patient-zero’ macaque by species (bonnet vs long-tailed vs rhesus macaques) for a given network-type. In each model, we included macaque group ID as a random effect, to account for intraspecific variation.

| Predictor | Model Coefficients |  |
| --- | --- | --- |
|  | Co-interaction networks | Grooming networks |
| (Intercept) | 1.20** | 1.19** |
| Sex (males vs females) | -0.03 | -0.05* |
| Rank percentile | 0.02 | 0.03* |
| Species (long-tailed vs bonnet) | -0.27 | -0.14 |
| Species (rhesus vs bonnet) | -0.23 | -0.65 |
| Species (long-tailed vs rhesus) | -0.04 | 0.50 |
| Strength (bonnet) | 0.49** | 0.25** |
| Strength (long-tailed) | 0.38** | 0.21** |
| Strength (rhesus) | 0.48** | 0.17** |
| Frequency of human-macaque interactions | 0.00 | 0.01 |
| Foraging on anthropogenic food | 0.02 | 0.01 |
| Strength (long-tailed vs bonnet) | -0.11* | -0.04 |
| Strength (rhesus vs bonnet) | -0.01 | -0.08* |
| Strength (long-tailed vs rhesus) | 0.10* | 0.04 |

\*p < 0.05; \*\*p < 0.01

For grooming networks, but not for co-interaction networks, we also found a significant effect of sex and dominance rank of the first-infected individual on mean outbreak sizes – zoonotic outbreak sizes were higher when first-infected macaques within grooming networks were females compared to males, and higher-ranking compared to lower-ranking individuals (Table 3). However, the magnitude of these effects were much lower than those of the strength centrality of first-infected macaques (Table 3). Finally, the overall anthropogenic exposure of first-infected macaques, i.e. their frequencies of interactions with humans and times spent foraging on human foods, had no impact on zoonotic outbreak sizes (Tables 2, 3).

## Discussion

We addressed a critical gap in our understanding of how zoonotic agents may spread and cause outbreaks among wildlife populations at HWIs. Adopting a comparative, network-based approach, we showed that zoonotic outbreaks among wild primate populations living in anthropogenic environments may reach higher outbreak sizes by spreading through animals’ joint social-ecological interactions with humans, than by spreading through their social interactions with conspecifics.

Zoonotic outbreak sizes were positively predicted by the centrality of the first-infected macaque within both their human co-interaction networks their grooming networks. By comparing the risk of zoonotic outbreaks posed by two different types of interactions, i.e. based on both interactions with conspecifics and with humans, we build on previous, network-based studies that have focused on modeling zoonotic outbreaks among wildlife populations through just interactions with conspecifics (e.g. European badgers, *Meles meles*: Rozins et al. 2018; chimpanzees: Rushmore et al. 2014; barbary macaques, *Macaca sylvanus*: Carne et al. 2017; interspecies comparative studies: Sah et al. 2018). Among the most widespread, ecologically flexible of all mammals outside of the family Rodentia, wild macaques may live in dense populations in a variety of anthropogenic environments (e.g. urban, agricultural, forest-fragmented habitats) throughout their geographic range, where they frequently interact with people. Thus, our finding that macaques’ tendencies to jointly interact with people make them especially highly vulnerable to zoonotic outbreaks has implications for our understanding of contemporary evolution and behavioral flexibility of wild animals living under increasing human impact. We previously speculated that such joint risk-taking may better enable wild primates to procure high-energy anthropogenic foods (Balasubramaniam et al. 2020b). Here, our findings suggest that the potential benefits of procuring such foods may be offset by high zoonotic risk. As such, our approaches in this study should encourage similar efforts on other wildlife populations that better distinguish between the (relative) effects of human-wildlife compared to wildlife-wildlife interactions on disease transmission and outbreaks among wildlife groups (e.g. elephants, *Loxodonta africana*, in agricultural fields: Chiyo et al. 2012; co-occurrence and space-use sharing of wild ungulates and livestock: VanderWaal et al. 2014b; human provisioning of birds and raccoons, *Procyon lotor*, in urban environments: Bradley & Altizer 2007).

For all three species, we found that the centrality of macaques within their co-interaction networks consistently led to higher zoonotic outbreak sizes compared to their centrality within grooming networks. In other words, the joint propensities for animals to aggregate around and interact with humans may lead to an even greater vulnerability of wildlife populations to zoonotic outbreaks than their interactions with conspecifics. This finding has major implications for “One Health” perspectives (Cunningham 2017; Zinsstag et al. 2011). To-date, research on disease transmission through wildlife populations has identified ‘superspreaders’ of pathogens that, in lieu of being more well-connected to other individuals and populations, may function as effective targets for disease control (e.g. vaccination, antimicrobial treatment: Drewe & Perkins 2015; Lloyd-Smith et al. 2005; Rushmore et al. 2014). Our findings suggest that macaques which are central in their human co-interaction networks may be especially effective targets, since these individuals may both function as intra-species superspreaders as well as pose a high risk of inter-species (humans-to-macaques, or vice-versa) disease spillover events since they inter-connect humans with whom they interact within and across time and space. Confirmation of this await future studies at HWIs that implement multi-modal networks that include pre-identified individual wildlife but also anthropogenic factors (individual humans, livestock, feral mammals) as nodes that are interlinked based on their shared space-use or social interactions (Silk et al. 2019). In particular, identifying points of wildlife-to-human disease spill-over are of utmost importance for preventing or controlling future pandemics like COVID-19 (Gryseels et al. 2020; Lappan et al. 2020).

We found cross-species differences in the extent to which co-interaction networks more strongly predicted zoonotic outbreak sizes compared to grooming networks. As predicted, rhesus macaques were the most vulnerable to zoonotic outbreaks through co-interaction networks, and the least vulnerable through their grooming networks. This highlights the importance of evaluating the relative effects of multiple (rather than single, as is often the case) single aspects of animal ecology on disease transmission. Rhesus macaques, more so than the other two macaque species, may preferentially engage in affiliative behaviors such as grooming with close kin (Thierry 2007); the resultant sub-grouping of individuals within their social networks may potentially function as ‘social bottlenecks’ to disease transmission in this species (Balasubramaniam et al. 2018; Griffin & Nunn 2012). Yet animals that show sub-divided social networks may nevertheless be vulnerable to outbreaks through other types of associations, and often in specific social-ecological contexts around human-provisioned food that may cause wild animals to aggregate together (Bradley and Altizer 2007) and co-interact with people (as we have shown).

Contrary to our predictions the effects of co-interaction networks on outbreak sizes in bonnet macaques were marginally greater (rather than lesser) than the effects of grooming networks, and were in fact within the range of rhesus macaques. One reason for this may be the spatial distribution of human-wildlife interactions in this population. Bonnet macaques are less geographically widespread and ecologically flexible compared to rhesus macaques. Although the bonnet macaques in our study experienced markedly lower frequencies of interactions with humans compared to rhesus macaques and long-tailed macaques (Krishna N. Balasubramaniam, Marty, Samartino, Sobrino, Gill, Ismail, et al. 2020), these interactions were highly geospatially restricted to within specific areas or ‘blocks’ within their home-range. It is likely that such spatially dense social-ecological associations with people, through increasing the connectivity of macaques within their co-interaction networks, leads to a considerable increase in the risk of zoonotic outbreaks despite their relatively lower overall frequencies of interactions with humans. More generally, this finding suggests that zoonotic agents may enter into and rapidly spread even through populations of less ecologically flexible wildlife that, despite interacting less frequently with humans, may congregate around anthropogenic factors within specific parts of their home-range (e.g., contexts of food provisioning: Marty et al. 2019b; crop-foraging: Chiyo et al. 2012; ecotourism activity: Carne et al. 2017). Aside from being the least ecologically flexible of the three species in this study, bonnet macaques are also the most vulnerable to human-impact, with many populations threatened by local extinction (Radhakrishna & Sinha 2011). Thus, the identification and treatment of potential superspreaders may be critical in this population.

Contrary to our prediction, long-tailed macaques showed no differences in zoonotic outbreak sizes across network-types. At least one explanation for this may be intra-specific variation, specifically between-group differences in their overall exposure to humans. We observed two groups of long-tailed macaques at a Hindu temple and popular tourist location within Kuala Lumpur, where the monkeys were exposed to dense human populations with whom they interacted highly frequently (Marty et al. 2019a). On the other hand, we observed two other groups in at a recreational park at the edge of the city bordering a fragmented forest area, where interactions with humans were comparatively less frequent (Marty et al. 2019a). Moreover, long-tailed macaques also showed marked differences in their grooming behavior across these locations as a response to interactions with humans (Marty et al. 2019a). This explanation seems to be supported by the separate groupings for the relationships between network centrality and outbreak sizes for long-tailed macaques, even for the same network-type (Figure 1). A more comprehensive assessment of the disease vulnerability of these populations would require within-species, cross-group comparisons.

Zoonotic outbreak sizes through macaques’ grooming networks were generally higher when the first-infected individuals were females compared to males, or when they were higher-ranking compared to lower-ranking individuals. Nevertheless, the effects of sex and dominance rank on zoonotic outbreaks were a lot weaker than the effects of individuals’ network centrality. In many wildlife species, animals’ sociodemographic attributes like their sex and dominance rank may influence their life-history, behavioral strategies, and adaptive responses to changing (anthropogenic) environments (Balasubramaniam et al. 2020b; Chiyo et al. 2012). It is therefore important to evaluate the potentially interactive effects of such factors with animals’ network connectedness on zoonotic outbreaks.

The consistently stronger effects of strength centrality compared to betweenness centrality or eigenvector centrality on outbreak sizes suggests that animals’ direct connections played a greater role in disease transmission than their secondary connections. This finding is largely consistent with previous epidemiological studies (Drewe & Perkins 2015), with some notable exceptions (e.g., betweenness as a stronger predictor of outbreaks across communities of humans (Funk et al. 2010) and chimpanzees (Rushmore et al. 2014). Such differences in the role of direct versus indirect connections in disease transmission may depend on the host population, network-type, or more global aspects of networks such as sub-grouping or *community modularity* (Griffin & Nunn 2012) or the *efficiency* of information transfer (Romano et al. 2018). Examining how these global aspects of macaques’ co-interaction networks and grooming networks may impact zoonotic transmission and outbreak sizes in these populations would be a critical next step.

Our results were independent of pathogen-specific transmissibility which, through influencing basic reproduction numbers (R_0_ values), may strongly impact zoonotic outbreaks. We chose to account for, rather than quantitatively evaluate, the effects of a suite of zoonotic respiratory pathogens of different transmissibility (e.g., influenza virus, measles virus, *Mycobacterium* spp., SARs-CoV-2), that typically spread through social interactions and are capable of causing disease in both humans and other primates (Rushmore et al. 2014; Sah et al. 2018). Pathogen transmissibility may interact with animal ecology in complicated ways to influence outbreak sizes. For instance, the effects of animal social interactions on zoonotic outbreaks may diminish for pathogens of exceptionally high transmissibility which may reach high outbreak sizes irrespective of social connections (Rushmore et al. 2014; Sah et al. 2018). Yet other studies have revealed that social interactions have stronger effects on outbreak sizes for pathogens of intermediate compared to low or high transmissibility (Rozins et al. 2018). Given the current lack of disease parameters on these macaque populations, our pathogen transmissibility values were also based on the human epidemiological literature (similar to other epidemiological studies on wildlife populations reviewed above). Inter-host and inter-pathogen differences would need to be considered in future studies that construct more sophisticated but system-specific epidemiological models.

## Acknowledgements

We thank the following organizations - the Himachal Pradesh Forest Department in India, Economic Planning Unit Malaysia, the Forestry Department of Peninsular Malaysia, the Department of Wildlife and National Parks Peninsular Malaysia, Tourism Selangor, and the Thenmala Forest and Wildlife Department - for their assistance through providing permission and logistical support to conduct research in India and Malaysia. Within these organizations, we are especially grateful to Drs. Lalith Mohan, Sandeep Rattan, Nadine Ruppert, Ahmad Ismail, Sahrul Anuar Mohd Shah, and Ullasa Kodandaramaiah, for their assistance and support. We grateful to research assistants Shelby Samartino, Mohammed Ismail, Taniya Gill, Alvaro Sobrino, Rajarshi Saha, Camille Luccisano, Eduardo Saczek, Silvia La Gala, Nur Atiqua Tahir, Rachael Hume, Kawaljit Kaur, Bidisha Chakraborty, Benjamin Sipes, Pooja Dongre, and Menno van Berkel for their involvement in data collection, processing, and storage in the field. The data for this study was collected as part of a human-primate Coupled Natural and Human Systems project supported by the U.S. National Science Foundation (Grant no. 1518555) awarded to PI McCowan.

## Author Contributions

K.N.B (first- and corresponding-author), under the supervision of E. A. and B.M., took the lead in in the study design, supervision of data collection, and the conductance of data analysis and manuscript writing. N. A. provided assistance with designing the study and writing the manuscript. B.A.B. and E.B.M. were involved in the formulation of field data collection procedures and manuscript writing. P.M., S.S.K., and M.A. were all involved in the designing and supervision of field-work (data collection), and participated in manuscript writing. E.A. and B.M. supervised the entire study.

