## Supplementary Tables and Figure for "Joint interactions with humans may pose a higher risk of zoonotic outbreaks than interactions with conspecifics among wildlife populations at human-wildlife interfaces"

Supplementary Table 1: Information on the study groups, subjects, and observation effort (table borrowed from our previous publication: Marty et al. 2019b)

| **Species** | **Group ID** | **Adult Males** | **Adult Females** | **Network nodes** | **Observation period** | **Focal Observation Hours (mean ± Std. dev.)** |
|---|---|---|---|---|---|---|
| Rhesus macaques (*Macaca mulatta*) | RM_G1 | 9 | 18 | 27 | Jun 2016 – Feb 2018 | 11.01 ± 3.14 |
| | RM_G2 | 7 | 17 | 24 | Jun 2016 – Feb 2018 | 14.46 ± 2.63 |
| | RM_G3 | 13 | 28 | 41 | Jun 2016 – Feb 2018 | 16.46 ± 5.74 |
| | RM_G4 | 14 | 45 | 59 | Jun 2016 – Feb 2018 | 8.50 ± 2.65 |
| Long-tailed macaques (*Macaca fascicularis*) | LM_G1 | 11 | 24 | 35 | Sep 2016 – Feb 2018 | 13.43 ± 4.29 |
| | LM_G2 | 7 | 12 | 19 | Sep 2016 – Feb 2018 | 13.5 ± 2.88 |
| | LM_G3 | 15 | 19 | 34 | Sep 2016 – Feb 2018 | 6.98 ± 1.47 |
| | LM_G4 | 5 | 19 | 24 | Sep 2016 – Feb 2018 | 12.16 ± 2.99 |
| Bonnet macaques (*Macaca radiata*) | BM_G1 | 26 | 22 | 48 | Jul 2017 – May 2018 | 11.78 ± 2.70 |
| | BM_G2 | 10 | 18 | 28 | Jul 2017 – May 2018 | 11.42 ± 1.41 |

Supplementary Table 2: GLMMs of network centrality by network-type for a given host-species, examining the effects of the attributes of the first-infected 'patient-zero' macaque. The models examined the effects of the attributes of the first-infected 'patient-zero' macaque, including an interaction between their network strength centrality and network-type, on mean outbreak sizes (proportion of infected macaques) calculated at the level of the individual across SIR model simulations run for pathogens of low to high transmissibility. We ran three model-sets, one for each host-species (A – C). Within each model-set, we ran three models, one for each of the three network centrality measures (strength, betweenness, and eigenvector). In each model, we included macaque animal ID (repeated measure across network-type) nested within group ID as random effects, to account for intraspecific variation. The model in bold indicates the best-fit model (AICc selection criterion of < 2 points from the next best-fit model) within each model set.

**(A) Bonnet macaques**

| **Model Number** | **Model ID** | **Predictor** | **B** | **SE** | **t** | **p** | **AICc** | **df** |
|---|---|---|---|---|---|---|---|---|
| **1** | **Interaction between strength centrality & network-type** | **(Intercept)** | **1.20** | **0.12** | **10.07** | **0.06** | **-699.51** | **143** |
| | | **Sex (males vs females)** | **-0.07** | **0.03** | **-2.16** | **0.03*** | | |
| | | **Rank percentile** | **0.01** | **0.03** | **0.41** | **0.68** | | |
| | | **Network (grooming vs co-interaction)** | **-0.10** | **0.03** | **-3.51** | **<0.01**** | | |
| | | **Network strength (co-interaction)** | **0.37** | **0.03** | **10.64** | **<0.01**** | | |
| | | **Frequency of interactions with humans** | **0.04** | **0.03** | **1.27** | **0.21** | | |
| | | **Foraging on anthropogenic food** | **0.04** | **0.03** | **1.46** | **0.15** | | |
| | | **Network strength (grooming vs co-interaction)** | **-0.19** | **0.06** | **-3.06** | **<0.01**** | | |
| 2 | | (Intercept) | 1.21 | 0.14 | 8.90 | 0.07 | -617.40 | 143 |

| | | | | | | | | |
|---|---|---|---|---|---|---|---|---|
| | Interaction between betweenness centrality & network-type | Sex (Males vs Females) | -0.19 | 0.04 | -4.90 | <0.01** | | |
| | | Rank Percentile | 0.07 | 0.04 | 1.71 | 0.09 | | |
| | | Network (grooming vs co-interaction) | -0.11 | 0.04 | -2.92 | <0.01** | | |
| | | Network betweenness (co-interaction) | 0.13 | 0.04 | 3.13 | <0.01** | | |
| | | Frequency of interactions with humans | 0.12 | 0.04 | 3.04 | <0.01** | | |
| | | Foraging on anthropogenic food | 0.04 | 0.04 | 0.94 | 0.35 | | |
| | | Network betweenness (grooming vs co-interaction) | -0.23 | 0.08 | -2.97 | <0.01** | | |
| 3 | Interaction between eigenvector centrality & network-type | (Intercept) | 1.19 | 0.09 | 12.76 | 0.05 | -679.59 | 143 |
| | | Sex (Males vs Females) | -0.06 | 0.04 | -1.68 | 0.09 | | |
| | | Rank Percentile | 0.01 | 0.03 | 0.33 | 0.74 | | |
| | | Network (grooming vs co-interaction) | -0.06 | 0.03 | -1.99 | 0.05* | | |
| | | Network eigenvector (co-interaction) | 0.36 | 0.04 | 9.13 | <0.01** | | |
| | | Frequency of interactions with humans | 0.06 | 0.03 | 1.96 | 0.05* | | |
| | | Foraging on anthropogenic food | 0.03 | 0.03 | 1.06 | 0.29 | | |
| | | Network eigenvector (grooming vs co-interaction) | -0.16 | 0.07 | -2.42 | 0.02* | | |

**(B) Long-tailed macaques**

| **Model Number** | **Model ID** | **Predictor** | **B** | **SE** | **t** | **P** | **AICc** | **df** |
|---|---|---|---|---|---|---|---|---|
| **1** | **Interaction between strength centrality & network-type** | **(Intercept)** | **0.95** | **0.21** | **4.55** | **0.02*** | **-905.20** | **213** |
| | | **Sex (males vs females)** | **-0.01** | **0.04** | **-0.31** | **0.76** | | |
| | | **Rank percentile** | **0.05** | **0.04** | **1.44** | **0.15** | | |
| | | **Network (grooming vs co-interaction)** | **-0.12** | **0.03** | **-3.57** | **<0.01**** | | |
| | | **Network strength (co-interaction)** | **0.22** | **0.04** | **5.92** | **<0.01**** | | |
| | | **Frequency of interactions with humans** | **0.03** | **0.04** | **0.87** | **0.39** | | |

| | | | | | | | | |
|---|---|---|---|---|---|---|---|---|
| | | **Foraging on anthropogenic food** | **0.01** | **0.04** | **0.34** | **0.74** | | |
| | | **Network strength (grooming vs co-interaction)** | **-0.06** | **0.07** | **-0.86** | **0.39** | | |
| 2 | Interaction between betweenness centrality & network-type | (Intercept) | 0.95 | 0.22 | 4.31 | 0.02* | -878.39 | 213 |
| | | Sex (Males vs Females) | -0.02 | 0.04 | -0.38 | 0.71 | | |
| | | Rank Percentile | 0.08 | 0.04 | 2.24 | 0.03* | | |
| | | Network (grooming vs co-interaction) | -0.11 | 0.04 | -3.00 | <0.01** | | |
| | | Network betweenness (co-interaction) | 0.08 | 0.04 | 2.32 | 0.02* | | |
| | | Frequency of interactions with humans | 0.06 | 0.04 | 1.47 | 0.14 | | |
| | | Foraging on anthropogenic food | 0.00 | 0.04 | -0.02 | 0.99 | | |
| | | Network betweenness (grooming vs co-interaction) | -0.07 | 0.07 | -0.96 | 0.34 | | |
| 3 | Interaction between eigenvector centrality & network-type | (Intercept) | 0.95 | 0.21 | 4.53 | 0.02* | -902.64 | 213 |
| | | Sex (Males vs Females) | -0.01 | 0.04 | -0.19 | 0.85 | | |
| | | Rank Percentile | 0.05 | 0.04 | 1.30 | 0.19 | | |
| | | Network (grooming vs co-interaction) | -0.09 | 0.03 | -2.69 | 0.01* | | |
| | | Network eigenvector (co-interaction) | 0.21 | 0.04 | 5.67 | <0.01** | | |
| | | Frequency of interactions with humans | 0.03 | 0.04 | 0.75 | 0.45 | | |
| | | Foraging on anthropogenic food | 0.03 | 0.04 | 0.69 | 0.49 | | |
| | | Network eigenvector (grooming vs co-interaction) | -0.01 | 0.08 | -0.10 | 0.92 | | |

**(C) Rhesus macaques**

| Model Number | Model ID | Predictor | B | SE | t | P | AICc | df |
|---|---|---|---|---|---|---|---|---|
| **1** | **Interaction between** | **(Intercept)** | **0.64** | **0.11** | **5.66** | **0.01*** | **-1638.94** | **291** |
| | | **Sex (males vs females)** | **-0.03** | **0.02** | **-1.43** | **0.15** | | |

| | | | | | | | | |
|---|---|---|---|---|---|---|---|---|
| | **strength centrality & network-type** | **Rank percentile** | **0.04** | **0.03** | **1.50** | **0.14** | | |
| | | **Network (grooming vs co-interaction)** | **-0.21** | **0.03** | **-6.50** | **<0.01**** | | |
| | | **Network strength (co-interaction)** | **0.91** | **0.04** | **21.69** | **<0.01**** | | |
| | | **Frequency of interactions with humans** | **0.02** | **0.04** | **0.46** | **0.65** | | |
| | | **Foraging on anthropogenic food** | **-0.04** | **0.04** | **-1.02** | **0.31** | | |
| | | **Network strength (grooming vs co-interaction)** | **-0.63** | **0.06** | **-9.96** | **<0.01**** | | |
| 2 | Interaction between betweenness centrality & network-type | (Intercept) | 0.83 | 0.12 | 6.79 | <0.01** | -1379.74 | 291 |
| | | Sex (Males vs Females) | -0.03 | 0.03 | -1.20 | 0.23 | | |
| | | Rank Percentile | 0.11 | 0.04 | 2.62 | 0.01* | | |
| | | Network (grooming vs co-interaction) | -0.35 | 0.03 | -11.24 | <0.01** | | |
| | | Network betweenness (co-interaction) | 0.37 | 0.06 | 6.09 | <0.01** | | |
| | | Frequency of interactions with humans | 0.13 | 0.06 | 2.14 | 0.03 | | |
| | | Foraging on anthropogenic food | -0.03 | 0.06 | -0.44 | 0.66 | | |
| | | Network betweenness (grooming vs co-interaction) | -0.33 | 0.08 | -3.95 | <0.01** | | |
| 3 | Interaction between eigenvector centrality & network-type | (Intercept) | 0.67 | 0.11 | 6.10 | 0.01* | -1595.15 | 291 |
| | | Sex (Males vs Females) | -0.03 | 0.02 | -1.80 | 0.07 | . | |
| | | Rank Percentile | 0.05 | 0.03 | 1.86 | 0.06 | | |
| | | Network (grooming vs co-interaction) | -0.18 | 0.03 | -6.60 | <0.01** | | |
| | | Network eigenvector (co-interaction) | 0.86 | 0.04 | 19.54 | <0.01** | | |
| | | Frequency of interactions with humans | 0.04 | 0.04 | 0.98 | 0.33 | | |
| | | Foraging on anthropogenic food | -0.04 | 0.04 | -0.97 | 0.33 | | |
| | | Network eigenvector (grooming vs co-interaction) | -0.67 | 0.06 | -10.69 | <0.01** | | |

**p < 0.01; *p < 0.05

Supplementary Table 3: GLMMs of network centrality by host-species for a given network-type, examining the effects of the attributes of the first-infected ‘patient-zero’ macaque. The models examined the effects of the attributes of the first-infected ‘patient-zero’ macaque, including an interaction between their network strength centrality and host-species, on mean outbreak sizes (proportion of infected macaques) calculated at the level of the individual across SIR model simulations run for pathogens of low to high transmissibility. We ran two model-sets, one for each network-type (A, B). Within each model-set, we ran three models, one for each of the three network centrality measures (strength, betweenness, and eigenvector). In each model, we included macaque group ID as a random effect, to account for intraspecific variation. The model in bold indicates the best-fit model (AICc selection criterion of < 2 points from the next best-fit model) within each model-set.

**(A) Co-interaction networks**

| Model Number | Model ID | Predictor | B | SE | t | p | AICc | df |
|---|---|---|---|---|---|---|---|---|
| **1** | **Interaction between strength centrality & species** | **(Intercept)** | **1.20** | **0.23** | **5.16** | **<0.01**** | **-1649.53** | **322** |
| | | **Sex (males vs females)** | **-0.03** | **0.02** | **-1.28** | **0.20** | | |
| | | **Rank percentile** | **0.02** | **0.02** | **0.84** | **0.40** | | |
| | | **Species (long-tailed vs bonnet)** | **-0.27** | **0.28** | **-0.94** | **0.38** | | |
| | | **Species (rhesus vs bonnet)** | **-0.23** | **0.28** | **-0.81** | **0.44** | | |
| | | **Strength (bonnet)** | **0.49** | **0.04** | **12.02** | **<0.01**** | | |
| | | **Frequency of human-macaque interactions** | **0.00** | **0.02** | **0.20** | **0.84** | | |
| | | **Foraging on anthropogenic food** | **0.02** | **0.02** | **1.04** | **0.30** | | |
| | | **Strength (long-tailed vs bonnet)** | **-0.11** | **0.06** | **-1.91** | **0.06** | | |
| | | **Strength (rhesus vs bonnet)** | **-0.01** | **0.05** | **-0.22** | **0.83** | | |

| | | | | | | | | |
|---|---|---|---|---|---|---|---|---|
| 2 | Interaction between betweenness centrality & species | (Intercept) | 1.29 | 0.24 | 5.39 | <0.01** | -1395.84 | 322 |
| | | Sex (males vs females) | 0.00 | 0.03 | -0.03 | 0.98 | | |
| | | Rank percentile | 0.08 | 0.03 | 2.73 | 0.01* | | |
| | | Species (long-tailed vs bonnet) | -0.43 | 0.29 | -1.44 | 0.19 | | |
| | | Species (rhesus vs bonnet) | -0.30 | 0.29 | -1.04 | 0.34 | | |
| | | Betweenness (bonnet) | 0.27 | 0.07 | 3.87 | <0.01** | | |
| | | Frequency of human-macaque interactions | 0.13 | 0.03 | 4.04 | 0.00 | | |
| | | Foraging on anthropogenic food | 0.03 | 0.03 | 1.03 | 0.30 | | |
| | | Betweenness (long-tailed vs bonnet) | -0.16 | 0.09 | -1.88 | 0.06 | | |
| | | Betweenness (rhesus vs bonnet) | -0.09 | 0.08 | -1.08 | 0.28 | | |
| 3 | Interaction between eigenvector centrality & species | (Intercept) | 1.20 | 0.23 | 5.26 | <0.01** | -1604.81 | 322 |
| | | Sex (males vs females) | -0.01 | 0.02 | -0.30 | 0.76 | | |
| | | Rank percentile | 0.02 | 0.02 | 1.18 | 0.24 | | |
| | | Species (long-tailed vs bonnet) | -0.28 | 0.28 | -1.02 | 0.34 | | |
| | | Species (rhesus vs bonnet) | -0.22 | 0.28 | -0.80 | 0.45 | | |
| | | Eigenvector (bonnet) | 0.46 | 0.04 | 10.83 | <0.01** | | |
| | | Frequency of human-macaque interactions | 0.04 | 0.02 | 1.71 | 0.09 | | |
| | | Foraging on anthropogenic food | 0.01 | 0.02 | 0.60 | 0.55 | | |
| | | Eigenvector (long-tailed vs bonnet) | -0.17 | 0.06 | -2.92 | <0.01** | | |
| | | Eigenvector (rhesus vs bonnet) | 0.00 | 0.05 | 0.02 | 0.99 | | |

**(B) Grooming networks**

| Model Number | Model ID | Predictor | B | SE | t | P | AICc | df |
|---|---|---|---|---|---|---|---|---|
| **1** | **Interaction between strength** | **(Intercept)** | **1.19** | **0.28** | **4.28** | **<0.01**** | **-2065.56** | **322** |
| | | **Sex (males vs females)** | **-0.05** | **0.01** | **-4.46** | **<0.01**** | | |
| | | **Rank percentile** | **0.03** | **0.01** | **3.06** | **<0.01**** | | |

| | | | | | | | | |
|---|---|---|---|---|---|---|---|---|
| | **centrality & species** | **Species (long-tailed vs bonnet)** | **-0.14** | **0.34** | **-0.42** | **0.69** | | |
| | | **Species (rhesus vs bonnet)** | **-0.65** | **0.34** | **-1.89** | **0.10** | | |
| | | **Strength (bonnet)** | **0.25** | **0.02** | **10.02** | **<0.01**** | | |
| | | **Frequency of human-macaque interactions** | **0.01** | **0.01** | **1.25** | **0.21** | | |
| | | **Foraging on anthropogenic food** | **0.01** | **0.01** | **0.71** | **0.48** | | |
| | | **Strength (long-tailed vs bonnet)** | **-0.04** | **0.03** | **-1.19** | **0.23** | | |
| | | **Strength (rhesus vs bonnet)** | **-0.08** | **0.03** | **-2.83** | **<0.01**** | | |
| 2 | Interaction between betweenness centrality & species | (Intercept) | 1.22 | 0.29 | 4.27 | <0.01** | -1868.46 | 322 |
| | | Sex (males vs females) | -0.15 | 0.01 | -10.50 | <0.01** | | |
| | | Rank percentile | 0.07 | 0.01 | 5.23 | <0.01** | | |
| | | Species (long-tailed vs bonnet) | -0.21 | 0.35 | -0.60 | 0.57 | | |
| | | Species (rhesus vs bonnet) | -0.66 | 0.35 | -1.89 | 0.10 | | |
| | | Betweenness (bonnet) | 0.00 | 0.03 | 0.09 | 0.93 | | |
| | | Frequency of human-macaque interactions | 0.02 | 0.01 | 1.53 | 0.13 | | |
| | | Foraging on anthropogenic food | -0.02 | 0.01 | -1.43 | 0.15 | | |
| | | Betweenness (long-tailed vs bonnet) | -0.01 | 0.04 | -0.13 | 0.89 | | |
| | | Betweenness (rhesus vs bonnet) | -0.02 | 0.04 | -0.43 | 0.67 | | |
| 3 | Interaction between eigenvector centrality & species | (Intercept) | 1.19 | 0.27 | 4.33 | <0.01** | -1998.10 | 322 |
| | | Sex (males vs females) | -0.08 | 0.01 | -5.75 | <0.01** | | |
| | | Rank percentile | 0.04 | 0.01 | 3.55 | <0.01** | | |
| | | Species (long-tailed vs bonnet) | -0.15 | 0.34 | -0.46 | 0.66 | | |
| | | Species (rhesus vs bonnet) | -0.63 | 0.34 | -1.89 | 0.10 | | |
| | | Eigenvector (bonnet) | 0.21 | 0.03 | 7.41 | <0.01** | | |
| | | Frequency of human-macaque interactions | 0.01 | 0.01 | 1.07 | 0.29 | | |
| | | Foraging on anthropogenic food | 0.00 | 0.01 | -0.09 | 0.93 | | |
| | | Eigenvector (long-tailed vs bonnet) | -0.04 | 0.03 | -1.10 | 0.27 | | |
| | | Eigenvector (rhesus vs bonnet) | -0.07 | 0.03 | -2.03 | 0.04 | | |

**p < 0.01; *p < 0.05

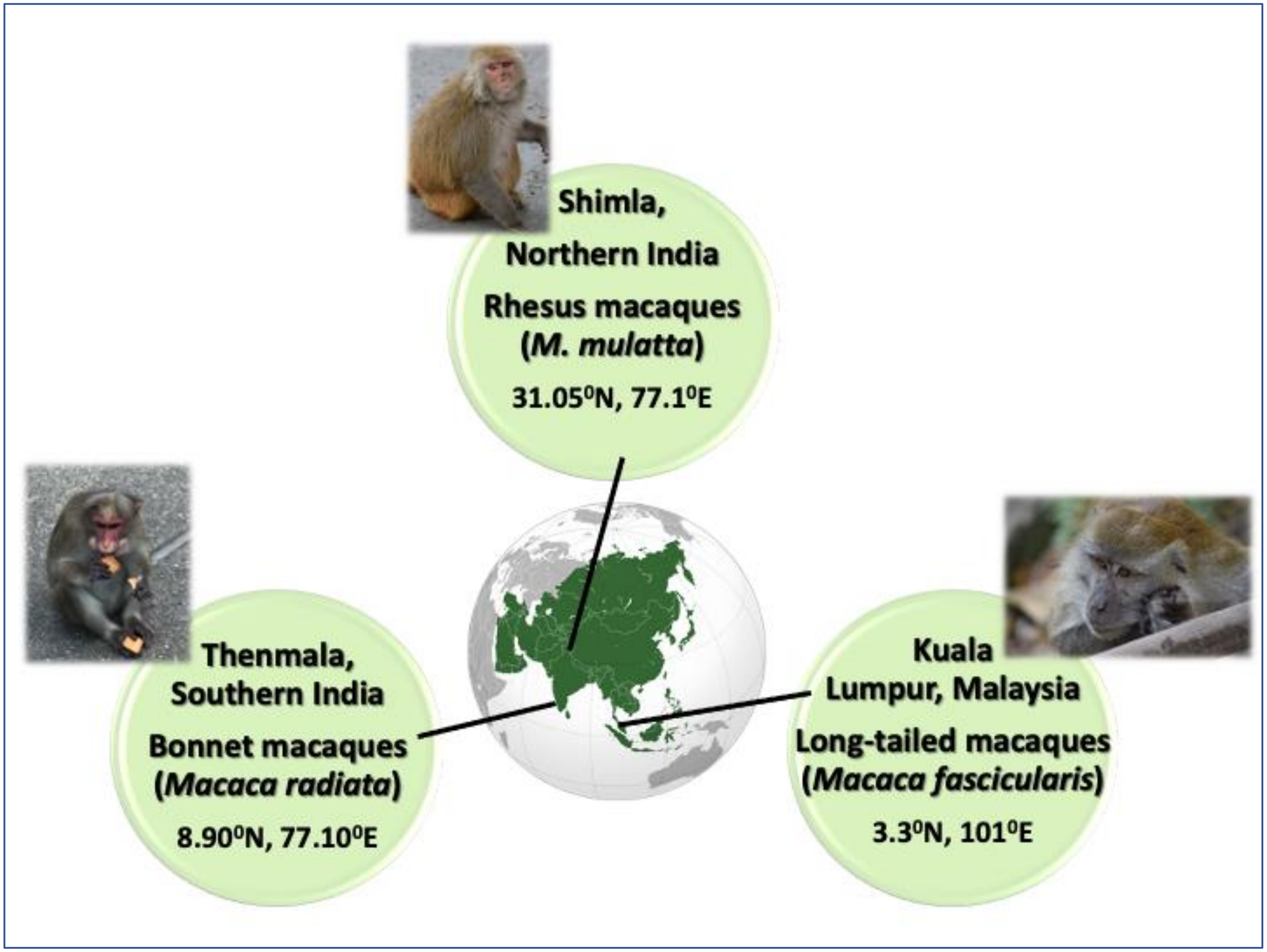


**Supplementary Figure 1: Macaque study species and locations at which the data were collected. Photo credits to K. N. B., S. S. K., and P. R. M.**
